# Enhanced visuomotor learning and generalization in expert surgeons

**DOI:** 10.1101/611012

**Authors:** Christopher L. Hewitson, David M. Kaplan

**Affiliations:** Department of Cognitive Science, Macquarie University, Sydney, Australia; Perception and Action Research Centre, Macquarie University, Sydney, Australia

## Abstract

Although human motor learning has been intensively studied for many decades, it remains unknown whether group differences are present in expert cohorts that must routinely cope with and learn new visuomotor mappings such as minimally invasive surgeons. Here we show that expert surgeons exhibit greater adaptation and generalization compared to naive controls in a standard visuomotor adaptation task. These findings run counter to a widespread background assumption in the field of motor learning that visuomotor adaptation performance should be largely uniform across the adult human population. Our findings also indicate that differences in basic visuomotor learning capacities, either innate or acquired, might be an important source of difficulty in learning and performing minimally invasive surgery. This information holds potential to guide surgical candidate selection or optimize training programs to address individual needs.

## Introduction

Human motor learning has been intensively studied for many decades [1, 2]. However, insufficient attention has been given to individual-or group-level differences in visuomotor learning. For example, it remains unknown whether group differences are present in expert cohorts that must routinely cope with and learn new visuomotor mappings such as minimally invasive surgeons.

Laparoscopic or minimally invasive surgery (MIS) is rapidly replacing traditional open surgery for many procedures due to its major benefits for patients over conventional open surgery including reductions in infection risks, recovery times, scarring, and overall hospital stays [3]. Despite these advantages, the task environment in MIS places high demands on surgeons, increasing the difficulty relative to open surgery for both initial learning [4] and ongoing performance [3,5,6]. Since laparoscopic instruments are controlled through small insertion points in the skin, instrument movements are often mirror-reversed and counter-intuitive (e.g., leftward hand motion produces rightward instrument tip motion, and vice versa). Because surgeons receive visual feedback indirectly via a laparoscopic camera that is in turn projected to a video display, rather than through direct observation, they must also contend with a range of visualization problems including absent depth information, variable magnification, and a restricted and frequently distorted (e.g., rotated) field of view. These factors, which are often subsumed under the general rubric of “challenges for hand-eye coordination” [7], also impose heavy computational demands on the brain and likely contribute to the significant increase in time to achieve proficiency in MIS compared to open surgery [8]; [9]

A related and potentially deeper explanation for why MIS is difficult to learn is that it requires complex sensorimotor transformations [10, 11]. Sensorimotor transformations involve the conversion of sensory inputs into appropriate motor commands [12], and are known to introduce errors even for simple goal-directed movements such as pointing to a visible target [13, 14, 15]. Moreover, because this sensory-to-motor mapping can and often does change during MIS such as when the laparoscopic camera rotates relative to the workspace or the fulcrum point of the instrument shifts, the same motor commands will not always lead to the same outcome. Consequently, surgeons must be particularly adept at learning new visuomotor mappings or transformations so they can maintain accurate motor performance during a procedure despite these changes. To appreciate the inherent challenges involved, one can imagine trying to use a computer mouse if the mapping between mouse and cursor movement changed frequently and unpredictably. Therefore, if MIS introduces challenges for learning appropriate sensorimotor transformations, this predicts that expert surgeons who have successfully learned to overcome these challenges, will perform better than naive controls in a standard visuomotor adaptation task [16, 17]. This is precisely the hypothesis we sought to test in this study.

Specifically, we predicted that expert minimally invasive surgeons would adapt more rapidly and more completely and would subsequently generalize their learning more widely than naive controls in a standard visuomotor adaptation task [17,18]. We found that expert surgeons both adapted faster and more completely and generalized to a greater extent across a range of novel target directions compared to controls. This study provides valuable insights into the improved visuomotor learning capacities among experts who draw on a vast reservoir of experience learning new visuomotor transformations. It provides evidence for the existence of group-level differences in visuomotor adaptation. Finally, it indicates that a standard visuomotor rotation task may have predictive validity for learning rates in training tasks specifically designed for MIS [19].

## Materials and Methods

### Participants

10 expert surgeons and 10 naive controls participated in the study. All were right-hand dominant with normal or corrected to normal vision. Experts (age 47±14 years, 9 males, 1 female) had all completed greater than 100 procedures (mean±SD = 2900±6000; range =>150<22000). Naïve controls (23±3 years, 4 males, 6 females) were university undergraduates with no prior medical experience, surgical training, or regular video game use (≤3 hours per week) [20–22]. All participants gave written informed consent to participate and the experimental protocols were approved by the [redacted for review] Human Research Ethics Committee (approval #5201833993962).

### Experimental Apparatus

A unimanual KINARM endpoint robot (BKIN Technologies, Kingston, Ontario, Canada) was utilized in the experiments for motion tracking and stimulus presentation. The KINARM has a single graspable manipulandum that permits unrestricted 2D arm movement in the horizontal plane. A projection-mirror system enables presentation of visual stimuli that appear in this same plane. Subjects received visual feedback about their hand position via a cursor (solid white circle, 2.5 mm diameter) controlled in real-time by moving the manipulandum. Mirror placement and an opaque apron attached just below the subject’s chin ensured that visual feedback from the real hand was not available for the duration of the experiment.

### Experimental Procedure

Subjects were instructed to perform fast and accurate reaching movements with the dominant (right) arm using cursor feedback, whenever it was available. Subjects performed reaches from a start target located at the center of the workspace to 11 different target locations 9 cm away from the start target and spaced 30° apart (Fig 1) The start target was a solid red circle (5 mm diameter), and each reach target was a solid green circle (5 mm diameter). The appearance of the reach target served as the go cue. Subjects were positioned so that the starting target was directly in front of their torso. Although these data are not reported here, a second workspace was also tested. In this experiment, the target array was translated rightward, which required subjects to extend their shoulder joint clockwise 45° to acquire the start target [23].

**Fig 1.**
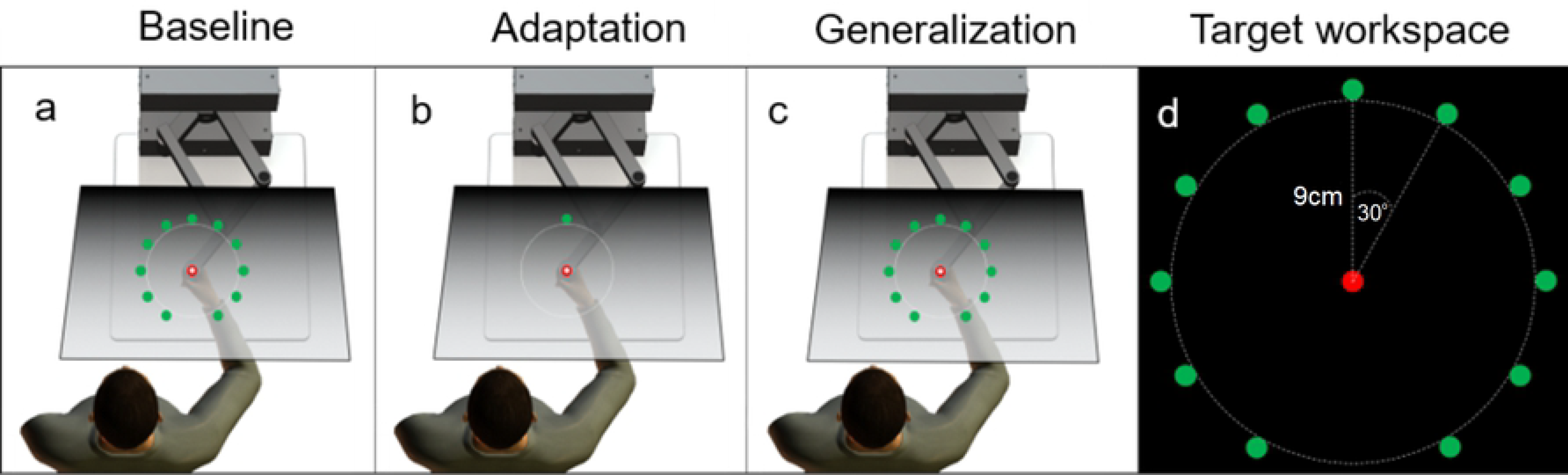
Experimental paradigm. (a) During the baseline block, subjects performed reaches to 1 of 11 targets with full visual feedback for 2/3 of the trials and no visual feedback for ⅓ of the trials, (b) During the adaptation block, subjects performed reaches to 1 target with visual feedback rotated at the endpoint of the movement, (c) During the generalization block, subjects performed reaches to 1 of 11 target (10 untrained target directions) without visual feedback, (d) Target array used in the experiment.

The experiment began with a familiarization block of 33 reach trials (3 per target in pseudorandom order) with veridical visual feedback provided throughout the reach. After the familiarization phase, subjects rested for 1 minute. The baseline block consisted of 198 reach trials (18 per target). For two-thirds of the reaches (132 trials), veridical cursor feedback was provided throughout the trial. For one-third (66 trials), visual feedback was withheld. During no-feedback trials, the cursor disappeared as soon as it left the start target. Visual feedback reappeared after the handle was positioned within a 1 cm diameter from the center of the start target. After the baseline phase, subjects rested for 1 minute. The adaptation block consisted of 110 reaches toward a single target positioned at 0° in the frontal plane. As the subject reached toward the target, cursor feedback was rotated about the start target by 30° (CW or CCW; counterbalanced between subjects). For the cursor to move directly toward the target, hand motion would need to be directed 30° opposite to the direction of the cursor rotation. For all reaches, no visual feedback was provided for the duration of the reach. For 90% of the trials, visual feedback was provided at the end of the movement for 150ms. For the remaining 10% of the trials, no visual feedback was provided during the reach or at the endpoint. The generalization block consisted of 66 reaches across 11 novel target directions (6 trials per direction) in pseudorandom order. Visual feedback was withheld for the entire outward reach movement and was only provided when the subject’s hand came within 1 cm of the start target during the return movement.

### Data Analysis and Models

Movement kinematics including hand position and velocity were recorded for all trials using BKIN’s Dexterit-E experimental control and data acquisition software (BKIN Technologies). Data was recorded at 200 Hz and logged in Dexterit-E. Custom scripts for data processing were written in MATLAB and data analysis was performed in JASP (0.9.2). A combined spatial-and velocity-based criterion was used to determine movement onset, movement offset, and corresponding reach endpoints [24, 25, 26]. Movement onset was defined as the first point in time at which the movement exceeded 5% of peak velocity after leaving the start target. Movement offset was similarly defined as the first point in time at which the movement dropped below 5% of peak velocity after a minimum reach of 9 cm from the start target in any radial direction. Reach endpoints were defined as the *x* and *y* values at movement offset. Reaction time (RT), total movement time (MT), peak velocity (PV), and endpoint error (EE; cursor position at movement offset - target position) were calculated for every trial across all experimental blocks and compared between groups and conditions via analysis of variance (ANOVA) and Welch T-tests. Learning rate was determined by fitting a power function to the data:

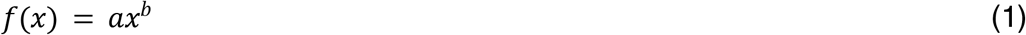

Because this assumes that learning is monotonic, which may not always be the case in visuomotor adaptation [27], data were also analyzed using repeated-measures ANOVA and Welch T-tests. Greenhouse-Geisser corrected values for ANOVAs are reported in case sphericity was violated. Generalization of learning to new target directions was compared between experts and controls using repeated-measures ANOVA and was quantified further by fitting subject data to a simple model in which generalization, *g*(*θ*), decreases as a Gaussian function of distance from the trained target direction [28, 23]:

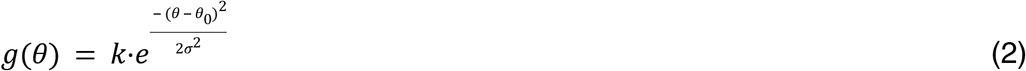

where *θ* is the trained target direction, *θ*_0_ is the peak or center of the generalization function (i.e., the target direction showing maximal adaptation), *k* is the magnitude or size of the peak (i.e., the amount of adaptation expressed in the maximal direction, and *σ* is the width. Goodness-of-fit was evaluated by calculating the coefficient of determination (*r*^2^).

## Results

To assess performance differences between groups, we measured reaction time (RT), total movement time (MT), peak velocity (PV), and endpoint error (EE; cursor position at movement offset - target position) for every trial across all experimental blocks. We began by investigating differences in reach performance between expert surgeons and controls prior to adaptation. During baseline reaches in which no visuomotor perturbation was imposed, experts and novices differed across a wide range movement parameters. Experts were more accurate (smaller endpoint error) and more precise (less variable) than controls (EE = 1.0±2.3° (SD) to the left of the target averaged across all target directions compared to EE = 3.9±5.7° to the left for controls; p<.001, Welch unpaired t-test) (Fig. 2). We also compared reaction time (RT = movement onset - go cue onset). Experts showed significantly (p<.001) faster reaction times (RT = 350±160ms) than controls (RT = 460±90ms). Next, we compared total movement time (MT = movement offset - movement onset). Experts were significantly (p<.001) quicker to complete reaches (MT =770±140ms) compared to controls (MT= 901±200ms). Interestingly, peak velocity was significantly (p<.001) greater for controls (PV = 15.0±0.7 cm/s) than for experts (PV = 14.0±0.6 cm/s).

**Fig 2.**
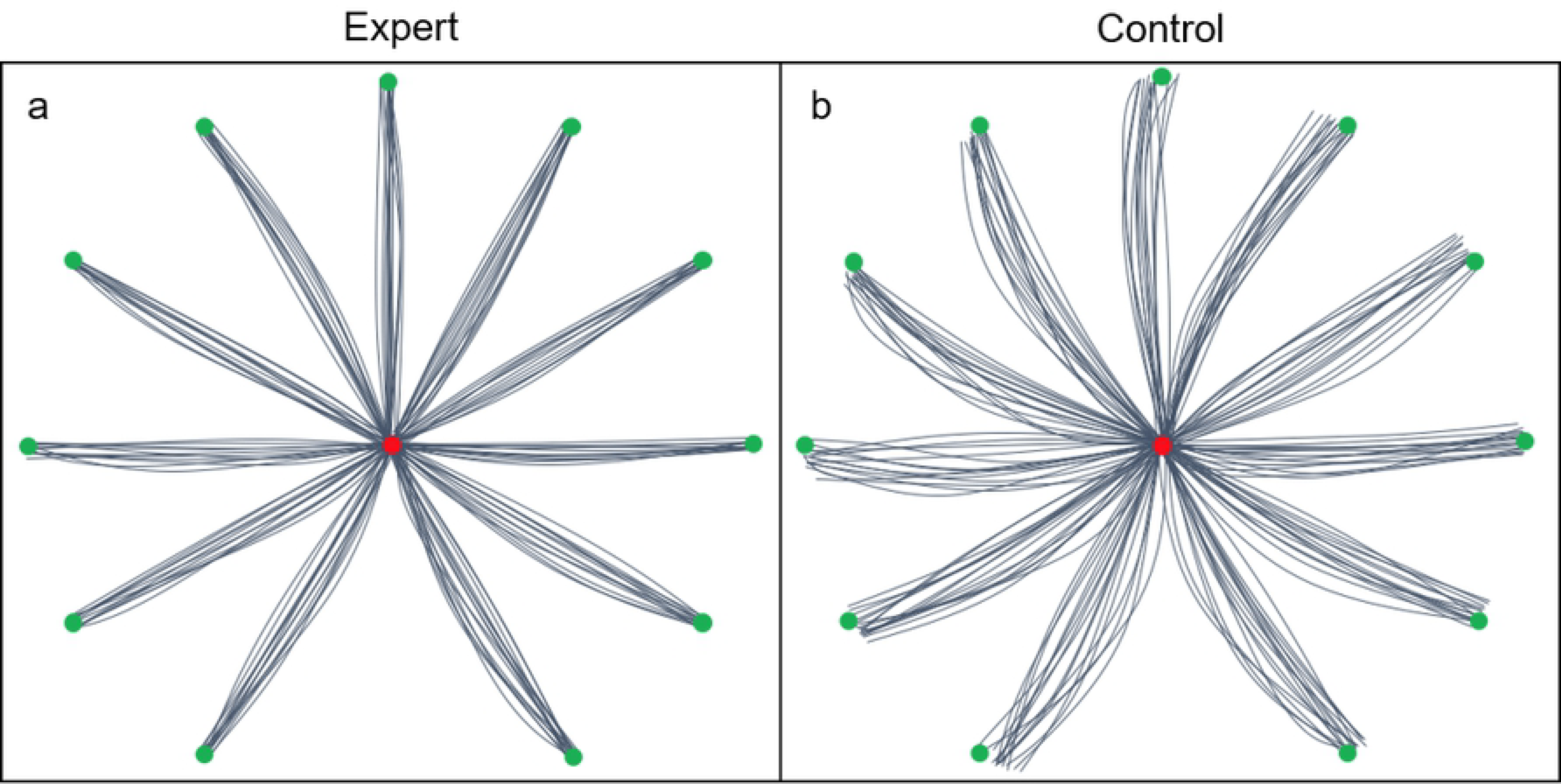
Baseline reaches for all target directions. (a) Representative expert, (b) Representative control.

Since this analysis collapsed across target directions and feedback conditions, we also performed ANOVAs for the within-subject factors of TARGET (11 target directions) and CONDITION (feedback vs no-feedback) and the between-subject factor of GROUP (experts vs controls) for all movement parameters. While there was a significant effect of GROUP, no GROUP × CONDITION × TARGET interactions were observed indicating no differences across target directions or feedback conditions.

Next, we tested whether performance during and after adaptation differed between experts and controls. We were specifically interested in whether there were group differences in the learning rate, which can be defined as the proportion of endpoint error that is corrected for in the subsequent movement. First, learning rates were estimated by fitting a decaying power function to the data (see Methods). Fitted values indicate that experts learned more rapidly than controls during the adaptation phase (*α* = 31.26, vs *α* = 26.48.8 respectively [Fig 3]). However, the coefficients of determination were relatively low (*r*^2^ = 0.472, *r*^2^= 0.377, respectively) so we also performed parametric comparisons. The within-subject factor of BIN and the between-subject factor of GROUP were compared via repeated-measures ANOVAs across 11 bins of 10 trials per bin. There was a significant within-subject effect of BIN (F_1,7.465_ = 59.6, p<.001, *ω*^2^ = 0.231; Greenhouse-Geisser corrected) as well as a significant between subject effect of GROUP (F_1,7.465_ = 269, p<.001, *ω*^2^ = 0.573; Greenhouse-Geisser corrected). These results indicate that while both groups adapted rapidly, experts did in fact adapt significantly faster than controls (Fig 3). In addition, averaged over the last two bins of trials during the adaptation block, experts adapted to the perturbation more completely than controls (29.6±2.0 [98.7%] compared to 28.5±4.3° [94.9%], p<.001; respectively, baseline adjusted). Reaction time and duration decreased over the course of the adaptation phase while peak velocity increased for both experts and controls (Fig 3). Although not our primary focus, there were significant within-subject effects of BIN as well as significant between subject effects of GROUP for all remaining movement parameters (Table 1).

**Table 1.** Baseline reach data for experts and controls.

|  | Endpoint error<br>(°) | Reaction time<br>(ms) | Movement time<br>(ms) | Peak velocity<br>(cm/s) |
| --- | --- | --- | --- | --- |
| Experts | 1.0±2.3° | 350±160 | 770±140 | 14.0±0.6 |
| Controls | 3.9±5.7 | 460±190 | 910±200 | 15.0±0.7 |

**Fig 3.**
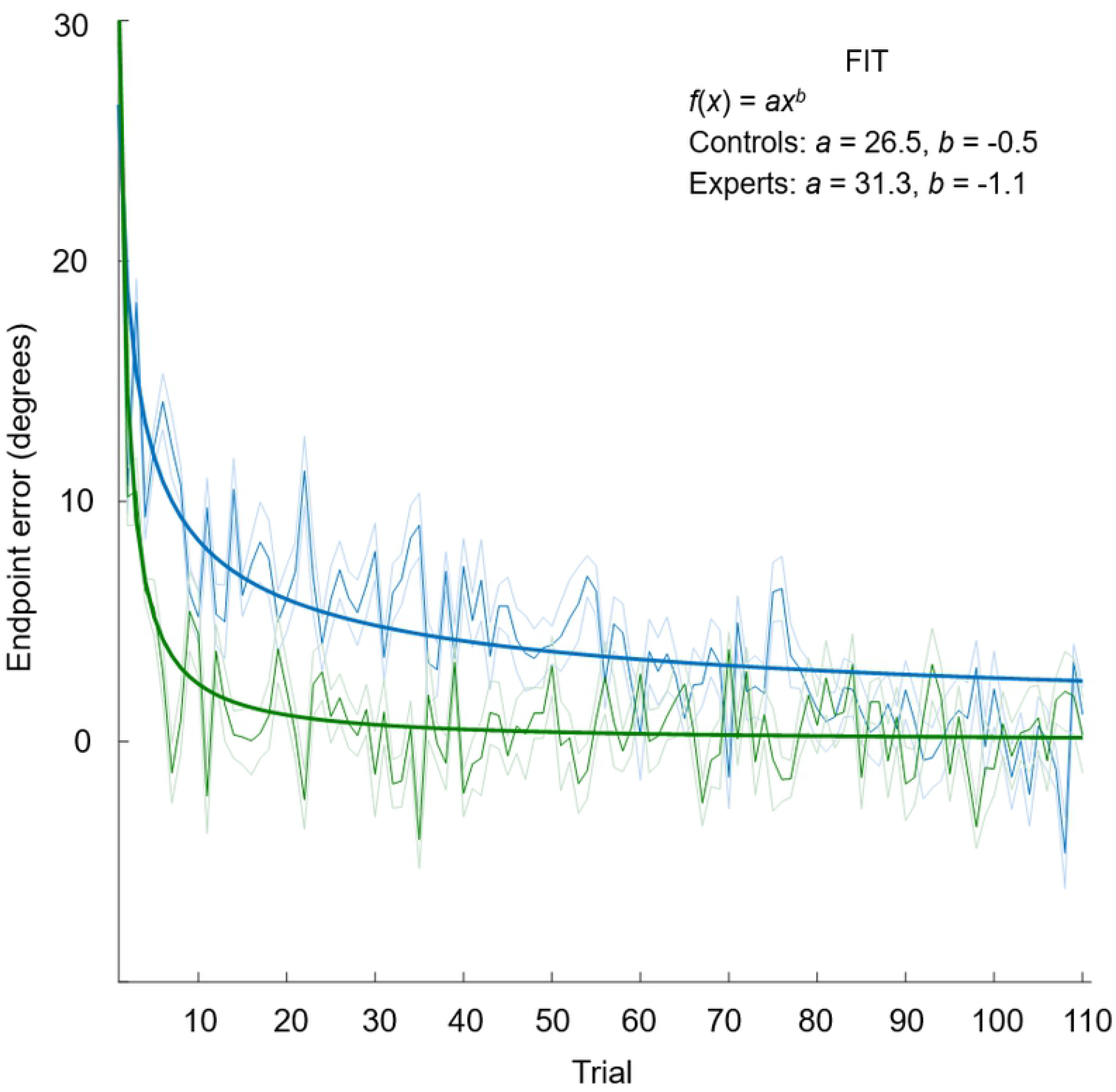
Learning rates for experts and controls. Green indicates experts and blue indicates controls. Thin solid lines show group means. Shaded region indicates s.e.m. Thick solid lines show model fits.

Finally, we investigated whether generalization of learning differed between the groups. Local generalization to new, untrained target directions decreased as a function of distance from the trained direction (up to ±90° from the trained target direction) for both experts and controls (Fig 4). The within-subject factor of TARGET and the between-subject factor of GROUP were compared via repeated-measures ANOVAs across the 11 targets with 6 trials per target. There was a significant within-subject effect of TARGET (F_1,8.674_ = 2518.2, p<.001, *ω*^2^ = 0.950; Greenhouse-Geisser corrected) as well as a significant between-subject effect of GROUP (F_1,8.674_ = 12276, p<.001, *ω*^2^ = 0.990). These results indicate that generalization varies with target direction, and that experts generalize significantly differently compared to controls.

**Figure 4.**
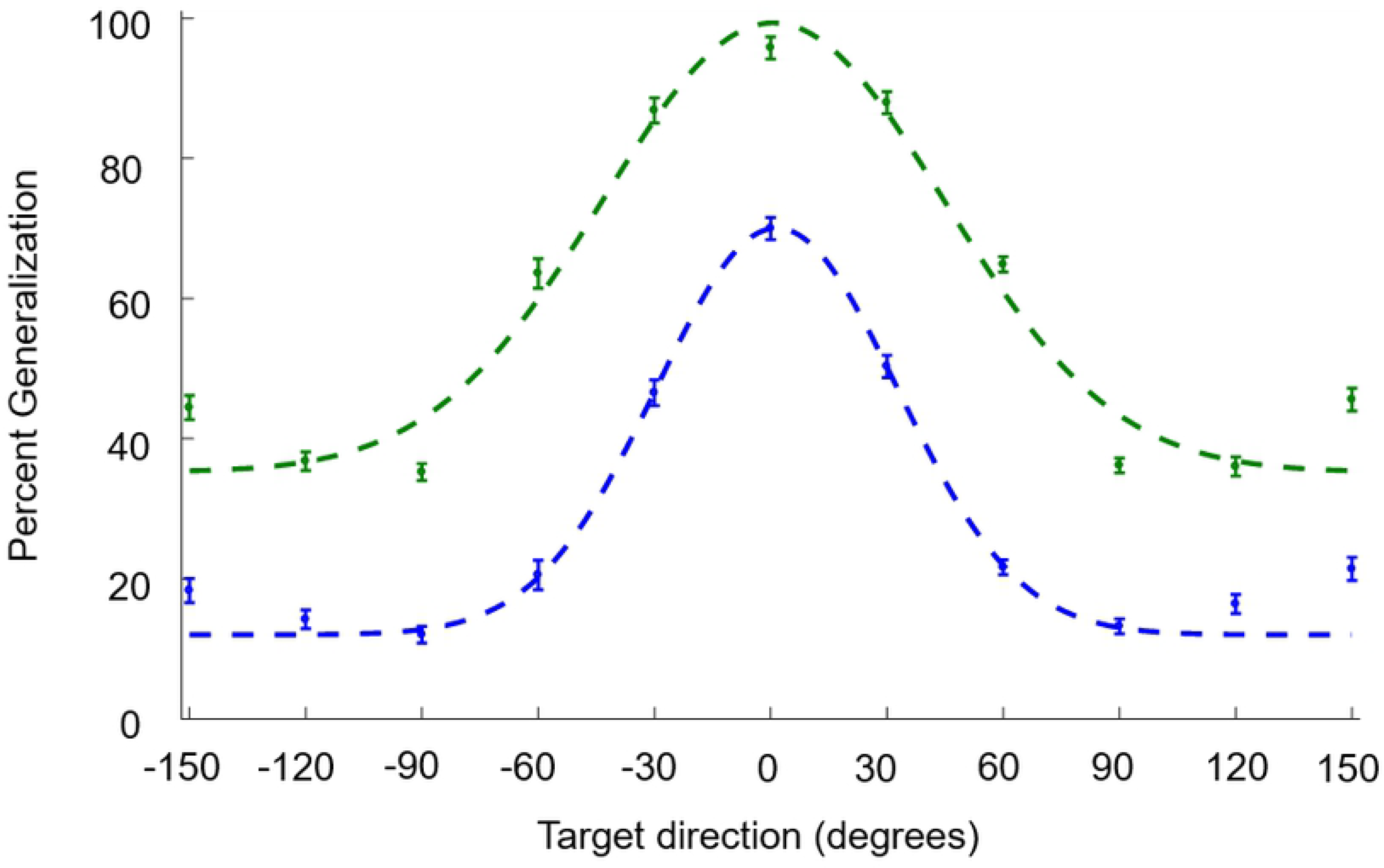
Generalization patterns for experts and controls. Green indicates experts and blue indicates controls. Error bars indicate s.e.m. Dashed lines indicate Gaussian model fits.

To further quantify the difference in generalization between experts and controls, endpoint data from targets were averaged across subjects and fit to a simple Gaussian model (see Methods) with four free parameters: center location, magnitude, width and vertical offset [23,28]. The fitted values for the magnitude and width parameters indicate that experts adapted to a far greater extent than controls (*k* = 97.1% vs *k* = 70.1%) and generalized their learning more broadly (*σ* = 43.9°, vs *σ* = 31.1°) than controls (*r*^2^ = 0.94, *r*^2^ = 0.96, respectively). Expressing generalization as a percentage of the actual adaptation amount rather than complete adaptation (100%) had little effect (98.4% vs 73.7%), since both groups almost completely adapted to the imposed visuomotor rotation by the end of the adaptation block.

## Discussion

In the current study, we found that expert minimally invasive surgeons exhibit enhanced visuomotor learning in every measure we tested. Specifically, experts adapted to the imposed visuomotor rotation more rapidly and more completely than controls. They also generalized their learning to new target directions more broadly and to a considerably greater extent.

What might explain these different learning patterns? One possibility is that attentional differences between surgeons and controls account for the observed differences in visuomotor learning performance. For example, there is some indication that extensive video game use can enhance visual selective attention [29]. Because of the coarse-grained similarities between video game playing and MIS (e.g., cursor control, complex task environment), it is possible that expert surgeons possess enhanced attentional capabilities, and this is what confers performance benefits in our visuomotor adaptation task. Despite its plausibility, specific features of our paradigm make this explanation unlikely. Even if the surgeons we tested have superior attentional capacities compared to controls, the task is not attentionally demanding since only a single reach target is presented per trial without the appearance of any distractors. Although attention has been shown to effect generalization of visuomotor learning [30, 31], all of these studies use concurrent attention-demanding secondary tasks during the adaptation period which differs substantially from the paradigm we employed.

Another, more probable explanation is that minimally invasive surgeons are exploiting additional resources such as explicit strategies or heuristics to adapt faster and more completely than controls. Although a long-standing view has been that visuomotor adaptation reflects incremental (i.e., trial-by-trial) error-based learning that occurs in an implicit and automatic manner, recent experimental and computational modelling work indicates that it often depends on the operation of multiple interacting learning processes [32,33]. For example, if a perturbation induces a large reaching error to the right side of the target, subjects experiencing this error signal might consciously choose to aim to the left side of the target on the next trial in order to compensate. One possibility is that our expert surgeons deployed similar strategies and did so more rapidly and effectively adapt than controls. Further studies will be required to tease apart the roles of these different visuomotor learning processes in expert surgeons.

Although visuomotor adaptation has been extensively studied [17, 2], few researchers have looked at learning differences across individuals or between specific groups. To date, there has been limited exploration of individual differences in visuomotor adaptation [34, 35], and very few studies have investigated adaptation differences between experts such as professional athletes and novices [36, 37]. This relative lack of attention may reflect a common background assumption in the field of motor learning that visuomotor adaptation performance should be largely uniform across the adult human population. Moreover, while expertise and expert performance have been thoroughly investigated from both psychological [38,39] and neuroscientific perspectives [40], surprisingly little attention has been given to surgeons as an informative expert cohort. The current study begins to close this important gap by showing that expert minimally invasive surgeons exhibit enhanced visuomotor learning in every measure we tested. Specifically, experts adapted to the imposed visuomotor rotation more rapidly and more completely than controls. They also generalized their learning to new target directions more broadly and to a considerably greater extent.

The findings reported here are striking because few studies have shown expert differences in visuomotor learning. In the most relevant study to date, Leukel and colleagues [36] explored how visuomotor learning might differ in expert handball players. Similar to our study, they reported lower movement variability (higher consistency) and greater accuracy in their expert group prior to learning, both widely considered to be hallmarks of expert performance [41]. But interestingly, they observed no learning rate differences in one experiment and a slower rate of adaptation in experts compared to novices in another experiment. One plausible explanation for the large discrepancy between their findings and ours is that handball players do not have to contend with or achieve mastery over changes in visuomotor mappings as do expert surgeons with extensive training and experience performing MIS.

Despite its major benefits for patients compared to open surgery, it is now widely recognized that minimally invasive surgery is inherently challenging to learn and can even be prohibitively difficult for some surgical residents such that they never reach proficiency [42,43,44]. Pinpointing the underlying sources of these learning curve differences remains an unanswered challenge. The findings reported here indicate that differences in visuomotor learning capacities, either innate or acquired, might be an important source of difficulty for learning to perform MIS. This information can be used to help guide surgical candidate selection or optimize training programs to address the specific needs of individual trainees. Our study also demonstrates that a general visuomotor learning paradigm from the field of motor neuroscience can reliably distinguish expert from non-expert in MIS. This result opens the door for the exploration of other common paradigms such as gain adaptation [18], which may shed valuable light on learning and expert performance in increasingly dominant approaches in surgical medicine such as robot-assisted minimally invasive surgery.

The authors declare no competing interests.

